# Fundamental Sex Differences in Cocaine-Induced Plasticity of D1R-and D2R-MSNs in the Mouse Nucleus Accumbens Core

**DOI:** 10.1101/2025.06.18.660420

**Authors:** Andrew D. Chapp, Hannah M. McMullan, Chau-Mi H. Phan, Pramit P. Jagtap, Paul G. Mermelstein

## Abstract

**BACKGROUND:** Previous studies have shown that cocaine-induced changes in nucleus accumbens shell (NAcSh) medium spiny neurons (MSNs) differ based on dopamine receptor subtype expression, the sex of the animal, and for females, phase of the estrous cycle. These findings highlight the need to account for both sex and estrous cycle when studying drug-mediated alterations in neurophysiology. Whether MSNs of the nucleus accumbens core (NAcC), which serve different aspects of addiction, will exhibit similar sex and estrous cycle effects with cocaine administration was investigated.

**METHODS:** Mice underwent a 5-day locomotor sensitization paradigm via daily cocaine administration (15 mg/kg, s.c.) followed by a 1-to 4-day drug-free abstinence period. We examined NAcC MSN excitability by obtaining *ex vivo* whole-cell recordings from differentially labeled dopamine D1-receptor expressing MSNs (D1R-MSNs) and dopamine D2-receptor expressing MSNs (D2R-MSNs) obtained from male mice or female mice that were either in estrus or diestrus.

**RESULTS:** In this genetic background of mice, both male and female mice sensitized to cocaine in a similar manner. In males, there were no cocaine-induced changes in D1R-MSN or D2R-MSN excitability, with D2R-MSNs exhibiting greater excitability. In saline-treated females, D1R-MSN excitability fluctuated across the estrous cycle with increased excitability during estrus. Following cocaine, estrous cycle-dependent D1R-MSN excitability was arrested, fixed at an intermediate value between estrus and diestrus when compared to saline controls. D2R-MSNs did not change either across the estrous cycle or following cocaine. When comparing MSN subtypes, in diestrus, D2R-MSNs were more excitable under saline conditions, but indistinguishable from D1R-MSNs following cocaine. In contrast, during estrus, D1R-and D2R-MSN excitability was similar in saline treated animals, but with cocaine, D2R-MSNs displayed heightened excitability.

**CONCLUSIONS:** There are fundamental sex differences in cocaine-induced changes to the excitability of D1R-MSNs in the NAcC. After cocaine exposure, female mice in diestrus exhibited a significant main effect change in MSN excitability, an inversion of what had previously been demonstrated in the NAcSh where no cocaine-induced changes were observed. These data suggest that there are distinct differences in the neuropharmacological effect of cocaine in males versus females that are shell and core specific.

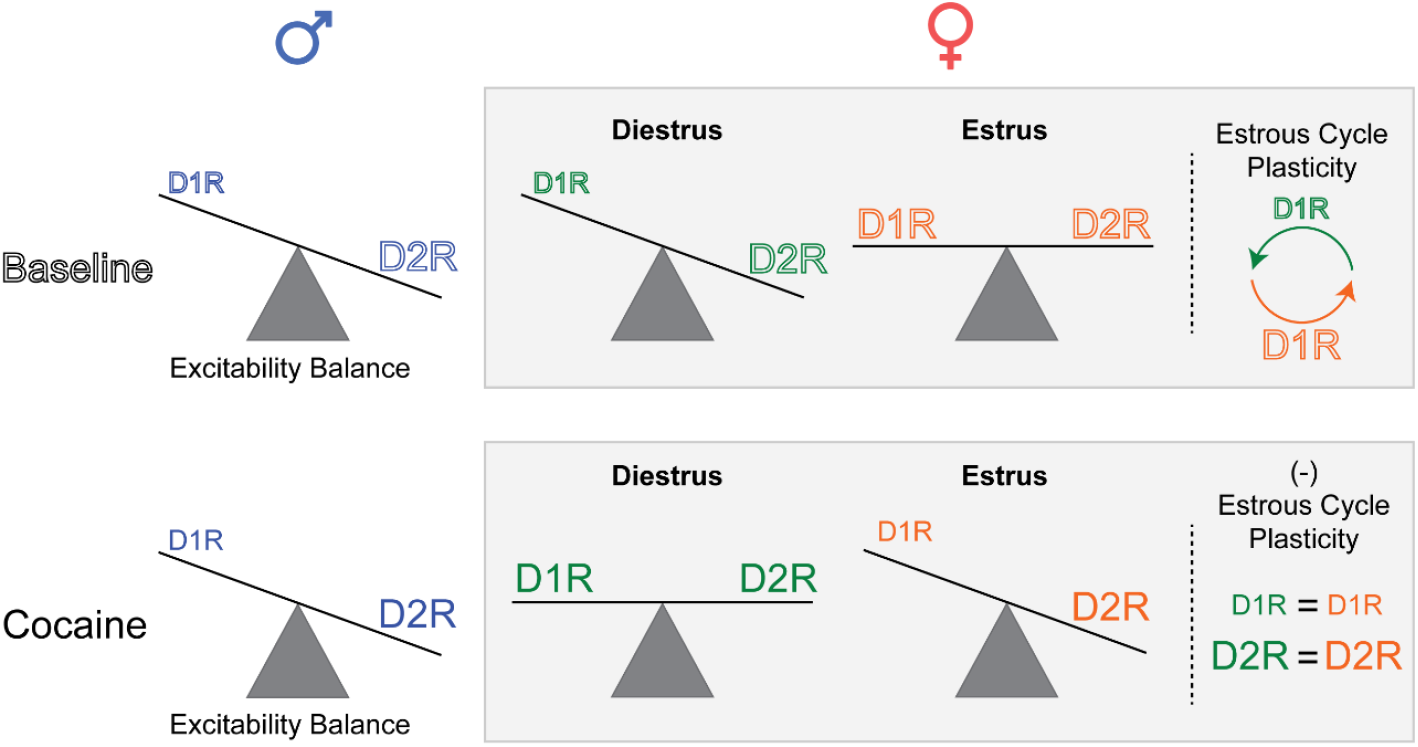

**HIGHLIGHTS:** There are sex-and estrous-cycle dependent changes to D1R-MSNs in the NAcC that are sensitive to cocaine exposure. In males, cocaine has no effect on D1R-or D2R-MSNs excitability. During the estrous cycle, D1R-MSNs exhibit increased excitability during estrus. This fluctuation is halted by cocaine, such that D1R-MSNs recorded in diestrus show increased excitability following cocaine exposure whereas female D1R-MSNs recorded in estrus have decreased excitability.

**PLAIN LANGUAGE SUMMARY:** The nucleus accumbens core (NAcC) is a brain region associated with regulating motivated behavior. The primary neuronal populations of the NAcC are dopamine D1 receptor expressing medium spiny neurons (D1R-MSNs) and dopamine D2 receptor expressing medium spiny neurons (D2R-MSNs). No studies exist which examine sex differences and estrous cycle effects in the NAcC following cocaine administration. Using *ex vivo* electrophysiology, we found inherent sex-and estrous-cycle differences in cocaine-induced changes in MSN neuroplasticity. D1R-MSN excitability was unaffected in males, increased in females recorded during the diestrus phase, and decreased in females recorded during estrus following cocaine exposure. This ran counter to estrous cycle effects under drug-naive conditions where D1R-MSN excitability was higher in estrus versus diestrus. The estrous cycle effects on D1R-MSNs were eliminated following cocaine administration. For both sexes, D2R-MSN excitability was not impacted following cocaine. These results highlight fundamental sex differences that might underpin differences in substance abuse.

## BACKGROUND

As of 2023, cocaine use disorder impacts 1.3 million Americans,^1^ and cocaine overdose deaths have increased nearly sixfold between 2011 and 2023.^2,3^ Cocaine exposure alters the intrinsic excitability of medium spiny neurons (MSNs) in the nucleus accumbens (NAc),^4–10^ with lasting plasticity in the NAc shell (NAcSh) and more transient alterations in the NAc core (NAcC).^5,11^ These brain regions are associated with regulating motivated behavior, and are thought to have differential roles in behavioral responses to psychostimulants such as cocaine. The NAcC is thought to regulate cue-associated drug seeking, while the NAcSh is linked to regulating behavioral responses such as locomotor sensitization.^12,13^ These drug-induced changes to MSN activity are proposed to contribute to the addictive properties of drugs of abuse.^14,15^ However, a fundamental limitation of prior work examining cocaine-induced plasticity in MSNs is that most studies were conducted using only male animals. Further, most studies that included females pooled data with males,^16–19^ and fewer still have tracked the estrous cycle. This is relevant because the nucleus accumbens exhibits estrous cycle-dependent plasticity thought to contribute to changes in motivated behavior across the estrous cycle.^20,21^

We have previously reported in the NAcSh that while cocaine decreases male D1R-MSNs excitability, it increases the excitability of D2R-MSNs in females in estrus and does not affect the excitability of either D1R-MSNs or D2R-MSNs of females in diestrus in the NAcSh.^22^ As the estrous cycle and sex hormones cause dynamic changes in the NAcC of rats^23–26^ in the absence of drugs, we hypothesized that estrous cycle plasticity may interact with cocaine-induced plasticity to differentially shape neuronal excitability in the NAcC. The excitability of MSNs in the NAcC is often the inverse of that observed in the NAcSh, with NAcC neurons increasing firing in early abstinence from psychostimulants and NAcSh decreasing firing in that same window (when D1R-and D2R-MSNs are not differentiated). This opens the question of whether the interactions of cocaine and sex hormones differs between the NAcC and the NAcSh.^27^

In this study, we expanded upon our previous research to assess whether there are sex differences in the NAcC MSN excitability both across the estrous cycle and following cocaine exposure. Using the same transgenic line of Drd1a-tdTomato and Drd2-eGFP mice as previously reported^22,28^ and powered to examine both sex and estrous cycle differences, we found both estrous cycle effects in MSN excitability and sex differences in response to cocaine distinct from those observed in the NAcSh.

## MATERIALS AND METHODS

### Animals

Animal procedures were performed at the University of Minnesota in facilities accredited by the Association of Assessment and Accreditation of Laboratory Animal Care and in accordance with protocols approved by the University of Minnesota Institutional Animal Care and Use Committee, as well as the principles outlined in the National Institutes of Health Guide for the Care and Use of Laboratory Animals. Male and female mice with a single copy of Drd1a-tdTomato^29^ and/or Drd2-eGFP^30^ bacterial artificial chromosome transgene were maintained on a C57BL/6J genetic background. Mice were originally obtained from the Rothwell lab (University of Minnesota)^28^ and bred onsite. Mice aged at least 8 weeks were used in all experiments, were group housed, and kept on a 14/10 hour light/dark cycle with food and water ad libitum. 62 animals were used in total, with 38 used for electrophysiology recordings (*n* = 12 males and *n* = 26 females).

### Psychomotor sensitization

All experiments were conducted between 10:00 AM and 6:00 PM, with house lights on at 6:00 AM and off at 8:00 PM. Animals were handled and habituated to locomotor chambers with subcutaneous saline injections 2 days prior to behavioral testing. On test days, animals were given either a subcutaneous injection of cocaine (15 mg/kg) daily for 5 days^5^ or an equivalent volume of 0.9% saline and placed immediately into the behavioral testing chamber (18” × 9”, with 8.5” tall walls) with light levels of 250 ±10 lx. Videos were recorded for 80 minutes using ANY-maze tracking software, and total distance travelled was analyzed offline.

### Chemicals

All chemicals were obtained from Sigma-Aldrich (St Louis, MO, USA), except cocaine hydrochloride and isoflurane which were obtained from Boynton Pharmacy (University of Minnesota, MN, USA).

### Whole-Cell Recordings

Mice (at least 8 weeks old) in early abstinence (1-4 days after the last behavioral day) were used for electrophysiology recordings. This time point was chosen based on prior work demonstrating that cocaine induces transient alterations in NAcC MSN excitability in male mice.^5,11^ Animals were sacrificed between 9:00 AM and 5:00 PM.

The estrous cycle was determined for females prior to anesthetization by vaginal cytology as previously described^18^. We chose to assess MSN excitability during diestrus and estrus in female mice as previous reports have shown that these windows have the greatest difference in neurophysiology.^26^ In mice, diestrus occurs during a trough in systemic estradiol, while estrus occurs shortly after the peak in estradiol concentrations. Mice were anesthetized with isoflurane (3% in O_2_) and decapitated. The brain was rapidly removed and chilled in ice-cold cutting solution containing: 228 mM sucrose, 2.5 mM KCl, 7.0 mM MgSO_4_, 1.0 mM NaH_2_PO_4_, 26 mM NaHCO_3_, 0.5 mM CaCl_2_, and 11 mM *d*-glucose with a pH 7.3 to 7.4 and continuously gassed with 95:5 O_2_:CO_2_ to maintain pH and pO_2_. A brain block containing the NAcC was prepared and affixed to the stage of a vibrating microtome (Leica VT 1200S; Leica). Sagittal sections of 240 µm thickness were cut, and the slices were transferred to a holding container of artificial cerebrospinal fluid maintained at 30 °C and gassed continuously with 95:5 O_2_:CO_2_ containing: 119 mM NaCl, 2.5 mM KCl, 1.3 mM MgSO_4_, 1.0 mM NaH_2_PO_4_, 26.2 mM NaHCO_3_, 2.5 mM CaCl_2_, 11 mM *d*-glucose, and 1.0 mM ascorbic acid (osmolality: 295-302 mosmol/L; pH 7.3-7.4)^7^ and allowed to recover for 1 hour. After recovery, slices were transferred to a glass-bottomed recording chamber and viewed through an upright microscope (Olympus) equipped with differential interference contrast optics, a SOLA SE light engine, fluorescence filters, an infrared (IR) filter, and a fluorescence/IR-sensitive video camera (Dage-MTI).

Slices were continuously perfused at ∼2 mL/min with room temperature artificial cerebrospinal fluid, gassed with 95:5 O_2_:CO_2_. Patch electrodes (5-10 MΩ open tip resistance) were pulled (Flaming/Brown P-97; Sutter Instrument) from borosilicate glass capillaries. Electrodes were filled with a solution containing 135 mM K-gluconate, 10 mM HEPES, 0.1 mM EGTA, 1.0 mM MgCl_2_, 1.0 mM NaCl, 2.0 mM Na_2_ATP, and 0.5 mM Na_2_GTP (osmolality: 280-285 mosmol/L; pH 7.3).^7^

D1R-MSNs and D2R-MSNs were identified under epifluorescence and IR-differential interference contrast based on soma morphology, tdTomato (D1-MSN) or EGFP (D2-MSN) reporter fluorescence, and a hyperpolarized membrane potential (-70 to-80 mV). Current-clamp recordings of membrane voltage were obtained using a Multiclamp 700B amplifier (Molecular Devices). Holding potentials were not corrected for the liquid junction potential. Once a GΩ seal was obtained, slight suction was applied to break into whole-cell configuration, and the cell was allowed to stabilize, which was determined by monitoring capacitance, membrane resistance, access resistance, and resting membrane potential (*V*_*m*_)7 using the membrane test function in pCLAMP acquisition software (Molecular Devices). Cells that met the following criteria were included in the analysis: action potential amplitude ≥ 50 mV from threshold to peak, resting *V*_*m*_ negative to-66 mV, and < 20% change in series resistance during the recording. The resting membrane potentials shown in Table 1 were recorded immediately after breaking into whole-cell mode. To measure NAcC MSN neuronal excitability, *V*_*m*_ was adjusted to-80 mV by continuous current injection as needed and a series of square-wave current injections was delivered from the baseline holding current in steps of +20 pA, each for a duration of 800 ms.

**Table 1.** Passive NAcSh MSN membrane properties.

| Experimental Condition | Capacitance (pF) | V<sub>m</sub> (mV) | R<sub>m</sub> (MΩ) |
| --- | --- | --- | --- |
| **Male D1R-MSN Saline (n=14)** | 67.79 ± 5.7 | - 75.50 ± 0.89 | 120.5 ± 8.8 |
| **Male D2R-MSN Saline (n=14)** | 67.14 ± 3.2 | - 73.64 ± 1.1 | 145.9 ± 11.7 |
| P value | 0.923 | 0.197 | 0.095 |
| **Male D1R-MSN Cocaine (n=15)** | 64.00 ± 2.6 | - 75.20 ± 1.1 | 118.8 ± 6.9 |
| **Male D2R-MSN Cocaine (n=14)** | 75.21 ± 5.1 | - 76.43 ± 1.2 | 119.8 ± 7.2 |
| P value | 0.066 | 0.467 | 0.922 |
| **Female Estrus D1R-MSN (n=14)** | 61.71 ± 12.3 | - 74.6 ± 4.4 | 159.7 ± 49.8 |
| **Female Estrus D2R-MSN (n=13)** | 66.5 ± 14.8 | - 71.7 ± 6.1 | 184.2 ± 72.5 |
| P value | 0.364 | 0.168 | 0.313 |
| **Female Estrus D1R-MSN Cocaine (n=16)** | 61.3 ± 10.7 | - 76.38 ± 2.8 | 117.4 ± 5.6 |
| **Female Estrus D2R-MSN Cocaine (n=16)** | 69.38 ± 17.7 | - 74.6 ± 5.6 | 122.4 ± 36.3 |
| P value | 0.130 | 0.260 | 0.695 |
| **Female Diestrus D1R-MSN Saline (n=17)** | 66.0 ± 2.6 | - 75.29 ± 0.94 | 117.8 ± 9.1 |
| **Female Diestrus D2R-MSN Saline (n=15)** | 74.7 ± 5.0 | - 74.07 ± 1.3 | 149.7 ± 14.4 |
| P value | 0.138 | 0.439 | 0.065 |
| **Female Diestrus D1R-MSN Cocaine (n=17)** | 78.4 ± 3.7 | - 76.3 ± 0.97 | 110.5 ± 8.5 |
| **Female Diestrus D2R-MSN Cocaine (n=16)** | 83.0 ± 5.5 | - 73.3 ± 1.3 | 96.4 ± 7.1 |
| P value | 0.482 | 0.077 | 0.218 |

Abbreviations: V<sub>m</sub>= Resting membrane potential, R<sub>m</sub>= membrane resistance.

### Statistical Analysis

Data values are reported as mean ± SEM. All statistical analyses were performed with a commercially available statistical package (GraphPad Prism, version 10.4.2). Probabilities < 5% were deemed significant a priori. Depending on the experiments, group means were compared using either a Student’s t-test, a Welch’s t-test (in the event there was a significant difference in variances), a 1-way or Welch’s 1-way repeated-measures analysis of variance (ANOVA), or 2-way repeated measures ANOVA. Where differences were found, Dunnett’s T3 post hoc tests were used for Welch’s ANOVAs and Fisher’s LSD post hoc tests were used for 2-way repeated measures ANOVA.

## RESULTS

We used a standard psychomotor sensitization protocol followed by a short, 1-to 4-day abstinence period prior to *ex vivo* whole-cell current clamp recordings of MSNs in the NAcC (Figure 1A)^5^ to examine cocaine sensitization and MSN excitability in male and female mice. Murine genetic background can produce large sex differences in cocaine-induced locomotor sensitization^31^. We therefore used a mouse line previously shown to have nearly identical male and female locomotor sensitization^31^ to ensure potential sex differences in NAcC D1R-MSN and D2R-MSN physiology were not attributable to behavioral differences. For electrophysiological recordings, parasagittal brain slices containing the nucleus accumbens were prepared from male mice or from female mice either in diestrus or estrus, when the largest differences in MSN physiology and behavior have been observed ^26,32^.

**Figure 1:**
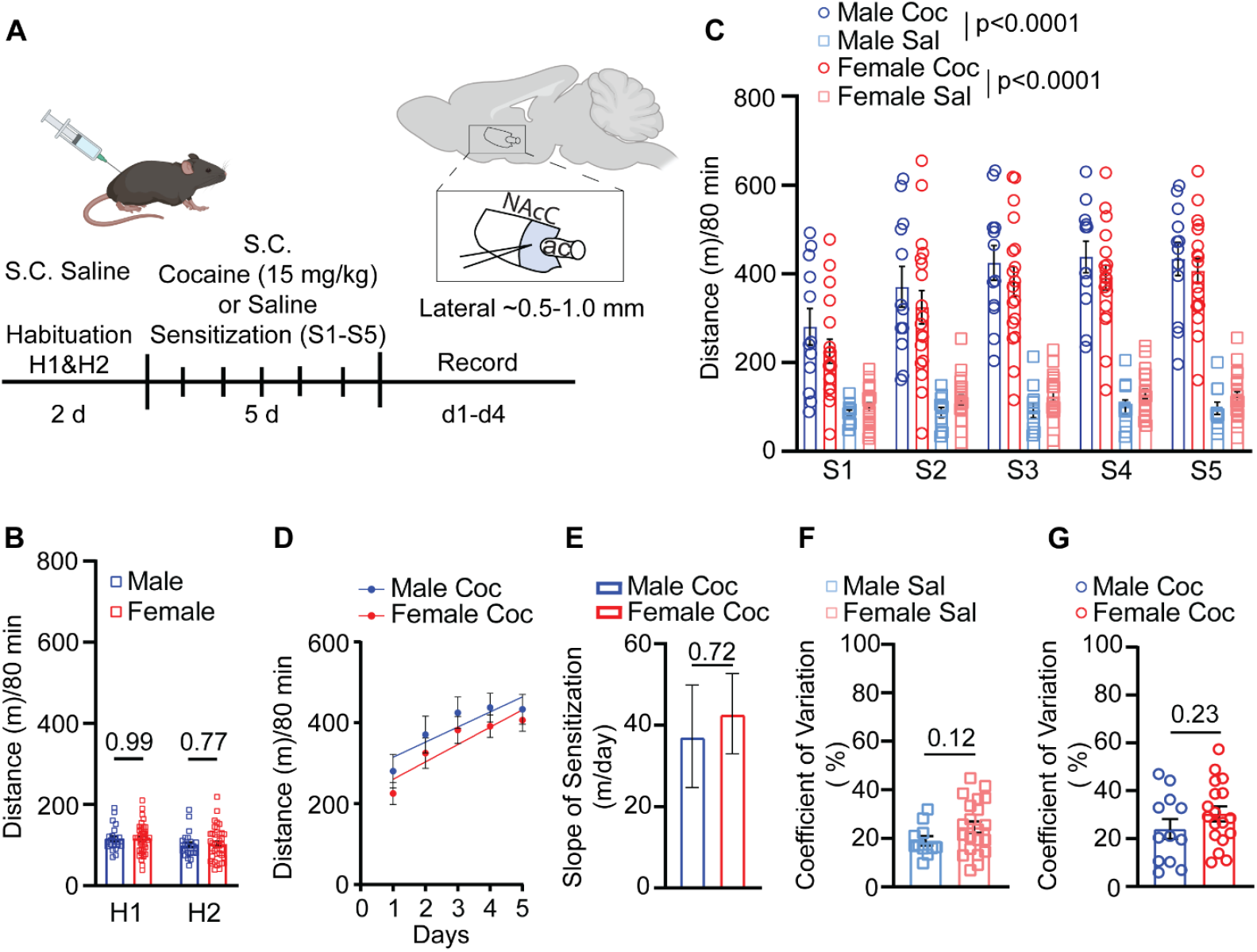
Cocaine psychomotor sensitization in male and female mice. **(A)** Experimental timeline and recording area of the NAcC highlighted in light blue. **(B)** Two-day saline habituation (H1 and H2) summary data of grouped male (*n* = 23) and female (*n* = 39) mice. **(C)** Psychomotor sensitization (cocaine: male *n* = 12, female *n* = 21; saline: male *n* = 11, female = *n* = 18). **(D)** Linear regression of cocaine psychomotor sensitization. **(E)** Slope of locomotor sensitization. **(F)** Coefficient of variation for saline-treated animals. **(G)** Coefficient of variation for cocaine-treated animals. Ac, anterior commissure; Coc, cocaine; NAcC, nucleus accumbens core; Sal, saline; S.C., subcutaneous.

All mice received saline during the 2-day habituation period (Figure 1B), after which they were subdivided into groups receiving either 15 mg/kg cocaine or saline for the next five days (Figure 1C). This dose has been used previously in our group for psychomotor sensitization and NAc neurophysiology experiments, and we have sought to maintain consistent comparisons for both behavioral sensitization and neuroplasticity studies^5,22^.

Cocaine increased the total distance travelled in both male (n =12) (2-way repeated measures ANOVA, F_1,21_ = 58.33, *p* < 0.0001) and female (n = 18) (2-way repeated measure ANOVA, F_1,37_ = 63.20, *p* < 0.0001) mice compared to their saline treated counterparts (*n* = 11 males, *n* = 21 females) (Figure 1C). Sensitization to cocaine was observed on day 2 of cocaine in both male (1-way repeated measures ANOVA, F_1.438,15.81_ = 10.03, *p* = 0.0031, Bonferroni’s post-test S1 v S2 *p* = 0.0034) and female mice (1-way repeated measure ANOVA, F_2.591,44.04_ = 16.76, Bonferroni post-test, *p* = 0.0008). We did not observe a difference in distance traveled between male and female cocaine-treated mice on any of the injection days (2-way repeated measure ANOVA, F_1,28_ = 0.9749, *p* < 0.3319). We further assessed whether there were any sex differences in cocaine sensitization by comparing the slope of total distance traveled on days 1 through 5 of cocaine treatment, representing the average increase in distance traveled per day within sex compared to the previous day (Figure 1D). We found no difference in the magnitude of sensitization between males and females (Figure 1E), confirming the minimal sex differences in cocaine-induced psychomotor sensitization in this strain of transgenic mouse^22^.

We also measured the coefficient of variation for each animal across the 5-day behavioral testing with respect to distance traveled and compared the coefficient across sexes. In contrast to previous reports, neither saline treated mice (Figure 1F) nor cocaine treated mice (Figure 1G) showed a difference in coefficient of variation between the sexes ^31^

### The Estrous Cycle and Early Abstinence from Cocaine Alters NAcC D1R-and D2R-MSN Excitability in a Sex-Dependent Manner

Following 1 to 4 days of abstinence, we investigated cocaine-induced changes to excitability in the NAcC among D1R-and D2R-MSNs. We recorded from 180 neurons in the medial portion of the NAcC. We observed lower D1R-MSN excitability relative to D2R-MSNs in both saline treated males (Figure 2A) (2-way repeated measures ANOVA, F_1,26_ =, *p* = 0.0280) and in cocaine treated males (2-way repeated measures ANOVA, F_1,27_ = 15.69, *p* = 0.0005) (Figure 2B). In contrast to what has previously been reported in the NAcSh, no change in excitability was observed in male D1R-MSNs following cocaine treatment (2-way repeated measures ANOVA, F_1,27_ = 0.3073, *p* = 0.5839) (Figure 2C, E). Similarly, no change in excitability was observed in male D2R-MSNs following cocaine treatment (2-way repeated measures ANOVA, F_1,26_ = 0.4273, *p* = 0.5191) (Figure 2D, F). This finding is consistent with the reports of He et al in male NAcC recordings from this strain.^11^

**Figure 2:**
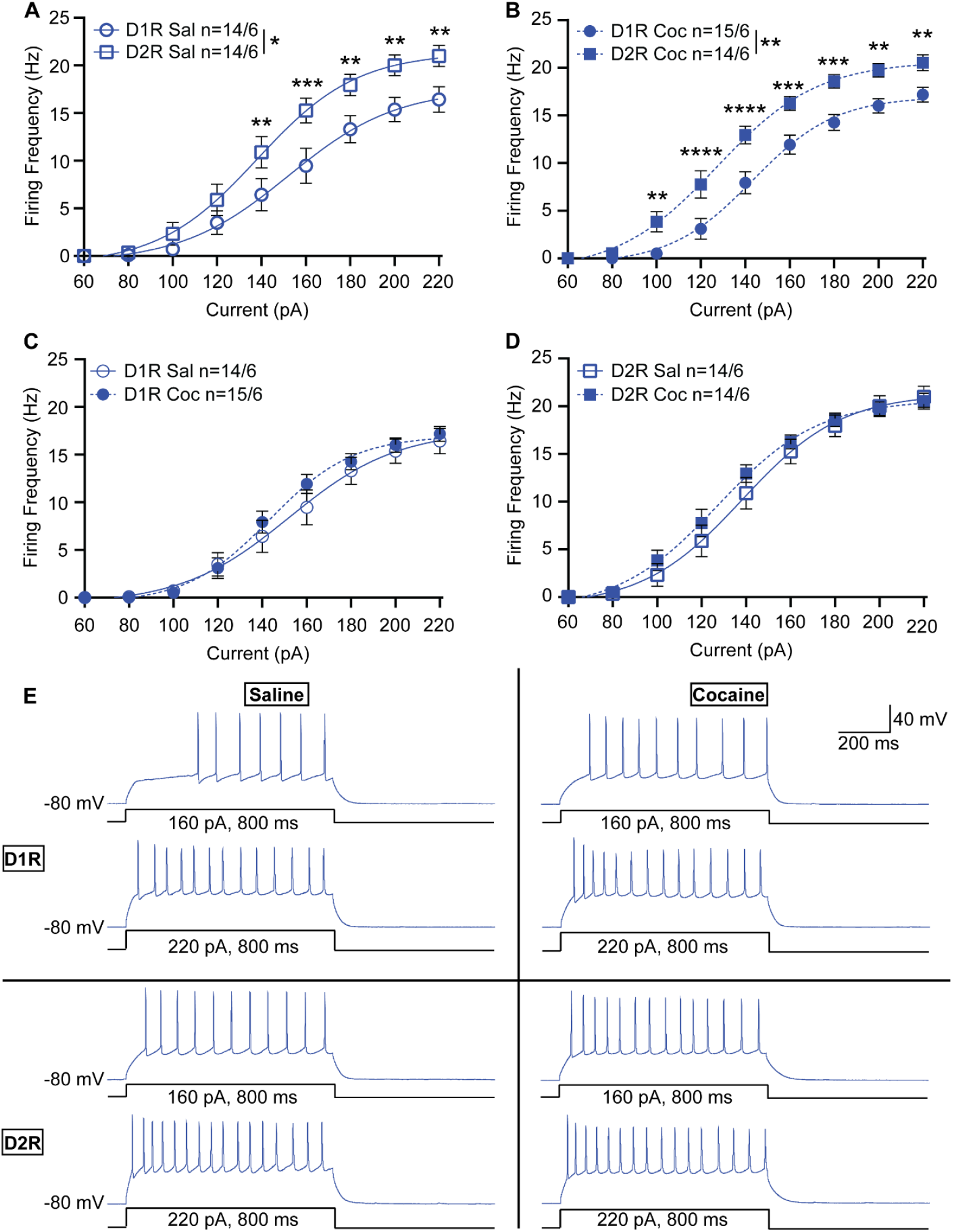
NAcC neuronal excitability in male mice. **(A)** Summary data for saline-treated male D1R-MSN vs. D2R-MSN neuronal excitability. **(B)** Summary data for cocaine-treated male D1R-MSN vs. D2R-MSN neuronal excitability. **(C)** Summary data for saline-vs. cocaine-treated male D1R-MSN neuronal excitability. **(D)** Summary data for cocaine-treated male D1R-MSN vs. D2R-MSN neuronal excitability. **(E)** Representative raw traces from saline-treated and cocaine-treated male D1R-MSN (top) and D2R-MSN (bottom) from the NAcC. The number of neurons and mice per group is denoted as n = x/y, where x is neurons and y are mice. Each group consisted of 6 mice. Coc, cocaine; D1R, D_1_ receptor; D2R, D_2_ receptor; Sal, saline. * p < 0.05, ** p < 0.01, *** p < 0.001.

Like males, female D1R-MSNs recorded in diestrus were less excitable than their D2R-MSN counterparts (2-way repeated measures ANOVA, F_1,30_ = 12.49, *p* = 0.0013) (Figure 3A). Interestingly, we observed no difference between D1R-and D2R-MSN excitability in cocaine-treated females recorded in diestrus (2-way repeated measures ANOVA, F_1,31_ = 0.6821, *p* = 0.4152) (Figure 3B). This was because in diestrus, D1R-MSNs of cocaine-injected female mice exhibited increased excitability compared to saline injected controls (2-way repeated measures ANOVA, F_1,32_ = 6.719, *p* = 0.0143) (Figure 3C).

**Figure 3:**
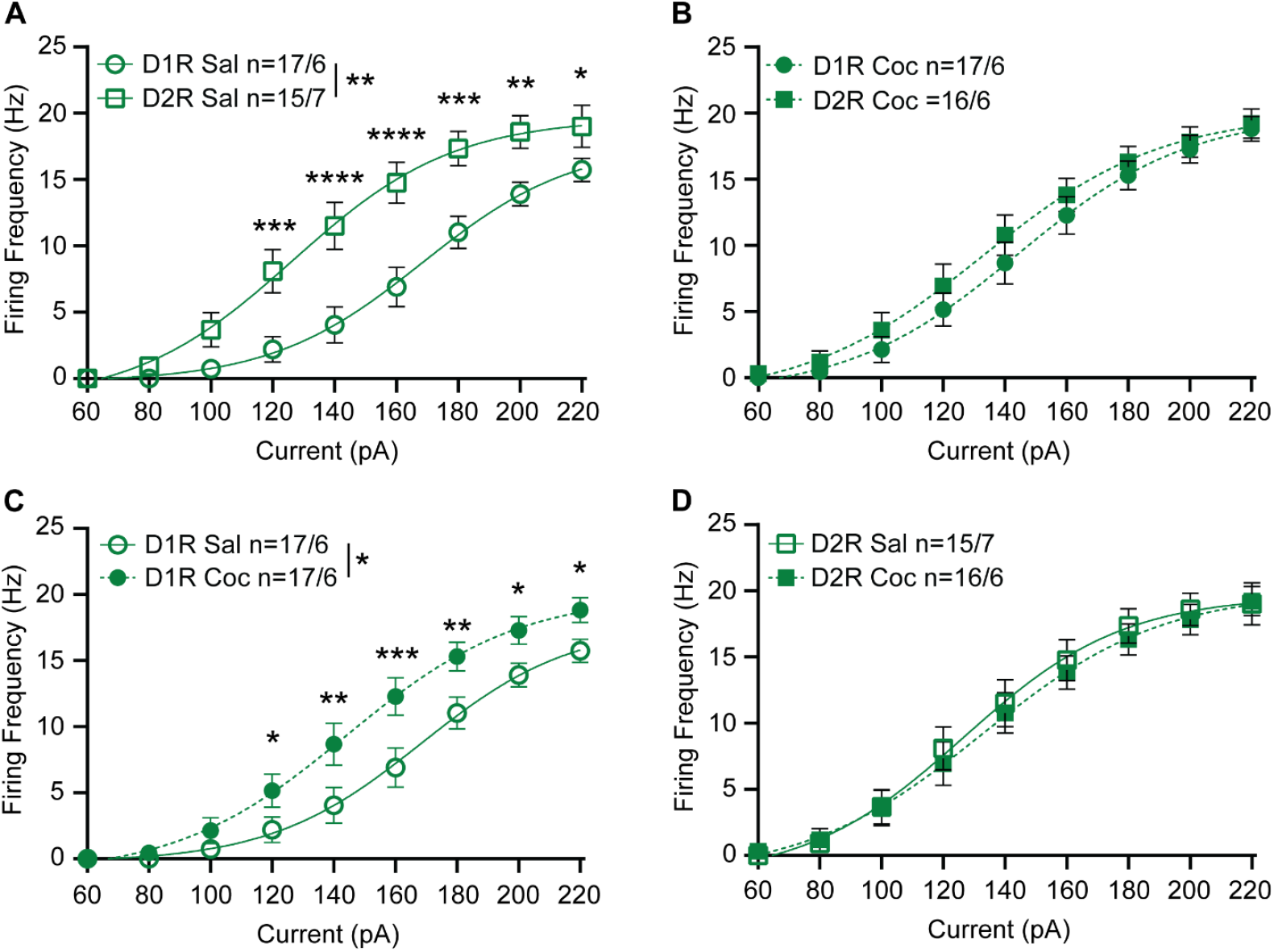
NAcC neuronal excitability in female mice recorded during diestrus. **(A)** Summary data for saline-treated female D1R-MSN vs. D2R-MSN neuronal excitability recorded during diestrus. **(B)** Summary data for cocaine-treated female D1R-MSN vs. D2R-MSN neuronal excitability recorded during diestrus. **(C)** Summary data for saline-vs. cocaine-treated female D1R-MSN neuronal excitability recorded during diestrus. **(D)** Summary data for cocaine-treated female D1R-MSN vs. D2R-MSN neuronal excitability recorded during diestrus. The number of neurons and mice per group is denoted as n = x/y, where x is neurons and y are mice. Each group consisted of 6 to 7 mice. Coc, cocaine; D1R, D_1_ receptor; D2R, D_2_ receptor; Sal, saline. * p < 0.05, ** p < 0.01, *** p < 0.001.

This finding is opposite to what has been observed in the NAcSh and in males, where D1R-MSNs exhibit decreased excitability following cocaine exposure.^22,33^ No change in excitability following cocaine exposure was observed in D2R-MSNs recorded from females in diestrus (2-way repeated measures ANOVA, F_1,29_ = 0.0823, *p* = 0.7762) (Figure 3D). In contrast to recordings from female mice in diestrus, we saw no difference between basal D1R-MSN and D2R-MSN excitability in females recorded in estrus (2-way repeated measures ANOVA, F_1,25_ = 0.1033, *p* = 0.751) (Figure 4A). Following cocaine exposure D1R-and D2R-MSNs exhibited differences in spiking activity for moderate levels of current injection (2-way repeated measures ANOVA, F_8,240_ = 2.949, *p* = 0.0037); significant differences observed at 120-160 pA step current injections (Figure 2B). These changes appeared to be primarily due to alternations in D1R-MSN activity following cocaine exposure. Following cocaine exposure, D1R-MSNs were significantly different from their untreated counterparts (2-way repeated measures ANOVA, F_8,224_ = 3.635, *p* = 0.0005); significant differences were observed at 120-180 pA injections (Figure. 4C). As with males and females recorded in diestrus, no change in D2R-MSN excitability following cocaine exposure was observed in females recorded in estrus (Figure 4D).

**Figure 4:**
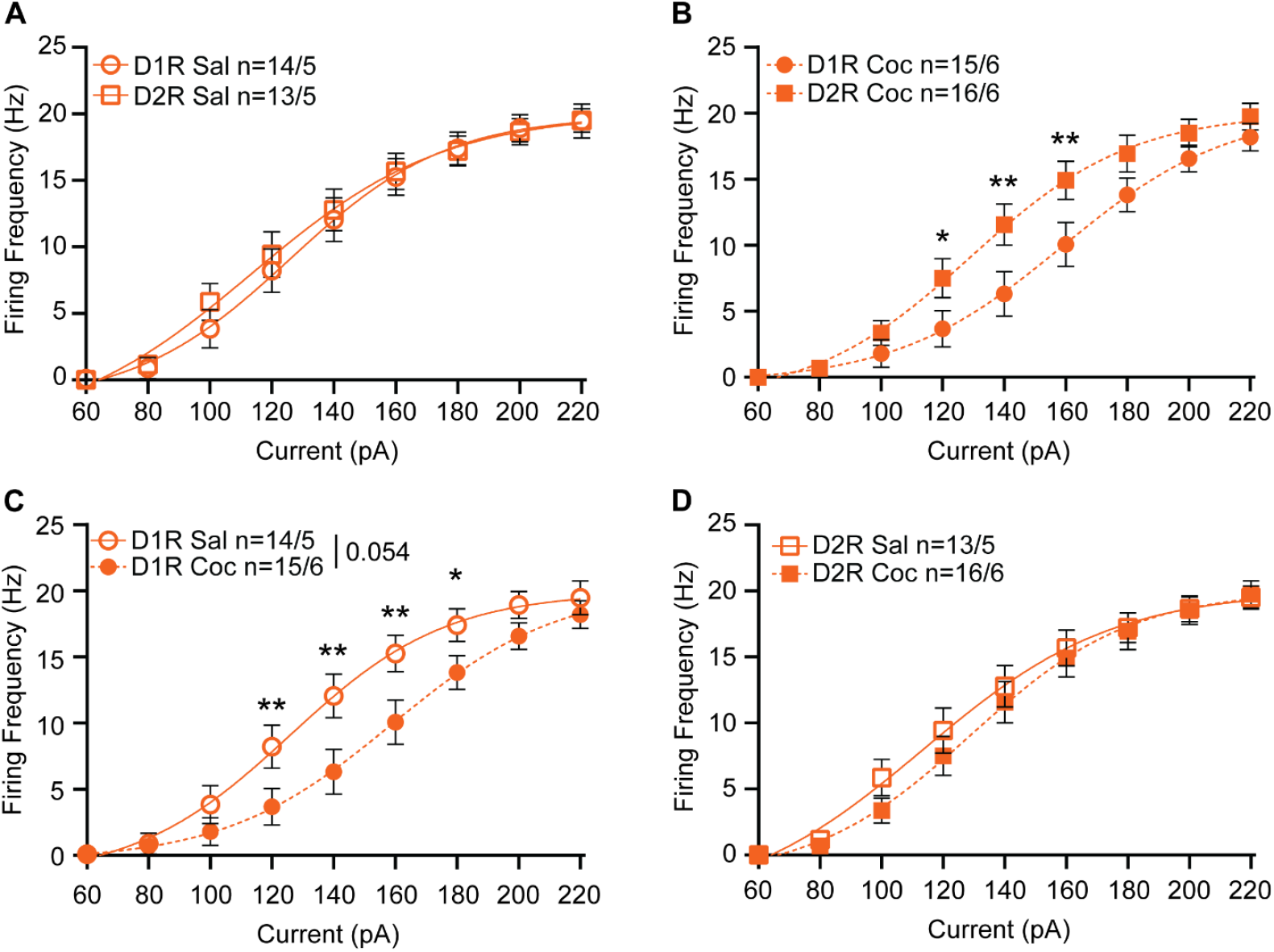
NAcC neuronal excitability in female mice recorded during estrus. **(A)** Summary data for saline-treated female D1R-MSN vs. D2R-MSN neuronal excitability recorded during estrus. **(B)** Summary data for cocaine-treated female D1R-MSN vs. D2R-MSN neuronal excitability recorded during estrus. **(C)** Summary data for saline-vs. cocaine-treated female D1R-MSN neuronal excitability recorded during estrus. **(D)** Summary data for cocaine-treated female D1R-MSN vs. D2R-MSN neuronal excitability recorded during estrus. The number of neurons and mice per group is denoted as n = x/y, where x is neurons and y are mice. Each group consisted of 5 to 6 mice. Coc, cocaine; D1R, D_1_ receptor; D2R, D_2_ receptor; Sal, saline. * p < 0.05, ** p < 0.01, *** p < 0.001.

Comparison of basal D1R-MSNs and D2R-MSNs excitability in females recorded during diestrus (Figure 3A) or estrus (Figure 4A) suggests that the estrous cycle modulates MSN intrinsic excitability. D1R-MSNs were more excitable in females recorded during estrus than in females recorded during diestrus (2-way repeated measures ANOVA, F_1,29_ = 14.07, *p* = 0.0008) (Figure 5A). This estrous cycle dependent effect was unique to basal D1R-MSN excitability, as no difference was observed between basal D2R-MSN excitability (2-way repeated measures ANOVA, F_1,26_ = 0.2521, *p* = 0.6199) (Figure 5B), cocaine-exposed D1R-MSN excitability (2-way repeated measures ANOVA, F_1,31_ = 0.6440, *p* = 0.4284) (Figure 5C), or cocaine-exposed D2R-MSN excitability (2-way repeated measures ANOVA, F_1,30_ = 0.0696, *p* = 0.7937) for females recorded in estrus or diestrus (Figure 5D). Thus, cocaine exposure abolished the estrous cycle-dependent plasticity in excitability of NAcC D1R-MSNs (Figure 5A,C).

**Figure 5:**
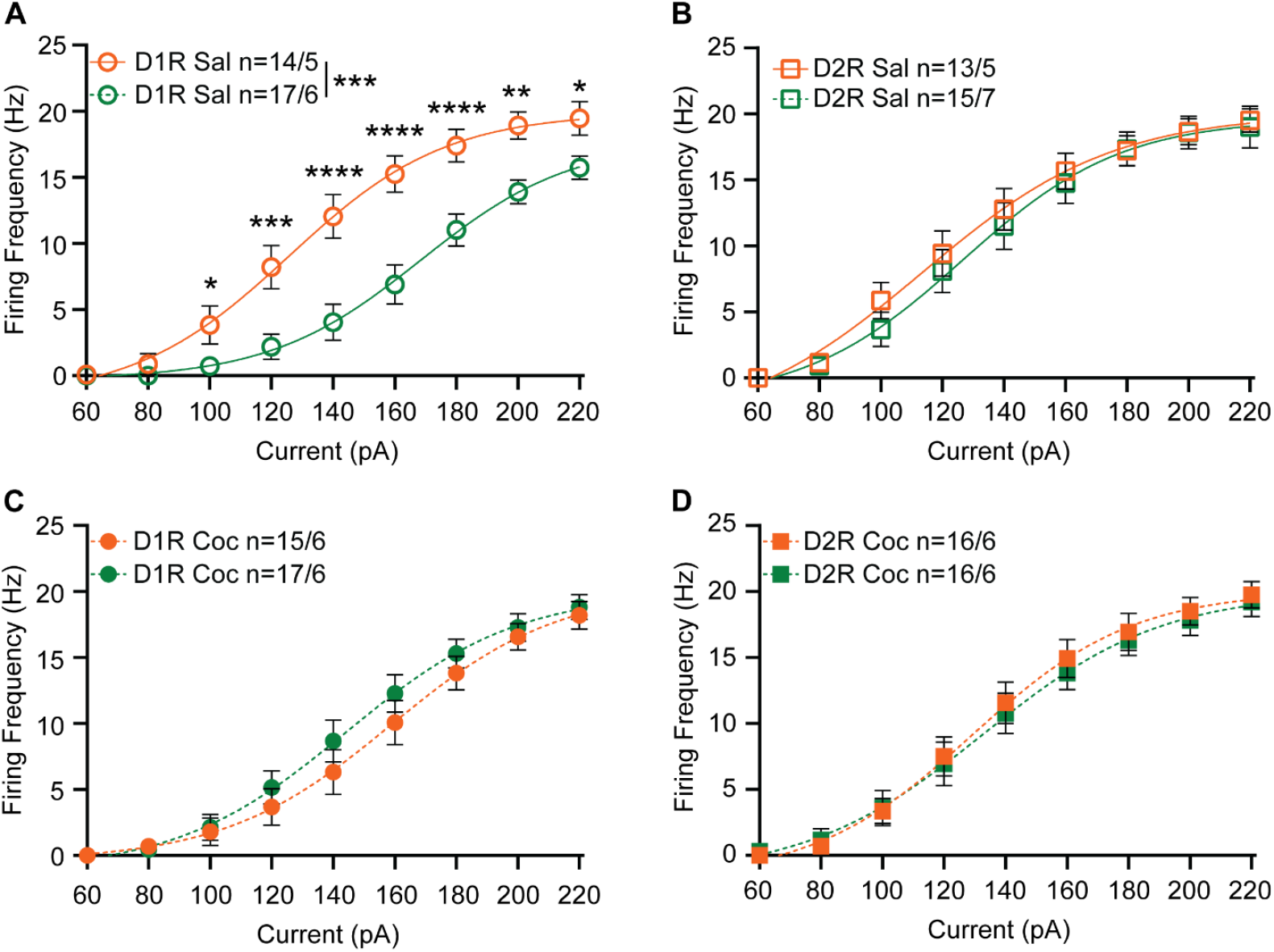
NAcC neuronal excitability in female mice recorded during diestrus (green) vs. recorded during estrus (orange). **(A)** Summary data for saline-treated female D1R-MSN neuronal excitability recorded during diestrus vs. recorded during estrus. **(B)** Summary data for saline-treated female D2R-MSN neuronal excitability recorded during diestrus vs. recorded during estrus. **(C)** Summary data for cocaine-treated female D1R-MSN neuronal excitability recorded during diestrus vs. recorded during estrus. **(D)** Summary data for cocaine-treated female D2R-MSN neuronal excitability recorded during diestrus vs. recorded during estrus. The number of neurons and mice per group is denoted as n = x/y, where x is neurons and y are mice. Each group consisted of 5 to 7 mice. Coc, cocaine; D1R, D_1_ receptor; D2R, D_2_ receptor; Sal, saline.

## DISCUSSION

This study expands upon our previous work to explore the effects of cocaine on the excitability of D1R-MSNs versus D2R-MSNs in the NAcC across sexes and with attention to the role of the estrous cycle on drug treatment.^22^ In agreement with what has been reported by He et al using a self-administration paradigm,^11^ there was no effect of cocaine on NAcC D1R-MSN or D2R-MSN excitability in males. Baseline MSN excitability in the NAcC mirrored our findings in the accumbens shell for males and diestrus females, where D2R-MSN excitability was greater than D1R-MSN excitability.^22^ This MSN subtype-specific difference in excitability was not observed in slices from females in estrus due to an increase in NAcC D1R-MSN excitability in estrus vs. diestrus, similar to our prior work in NAcSh.^22^

D1R-MSNs and D2R-MSNs are thought to have differential roles. For example, optogenetic stimulation of striatal D1R-MSNs in male mice has rewarding effects whereas identical stimulation of D2R-MSNs is aversive.^34,35^ While studies in male animals have primarily implicated a role for D1R-MSNs in the rewarding effects of cocaine, studies in female animals have suggested that an altered balance of D1R-MSN and D2R-MSN activity may be the relevant variable modulating drug reward.^17,36^ Consistent with this latter model, our previous work showed there are sex differences following cocaine exposure in the NAcSh, with a decrease in D1R-MSN activity in males and an increase in D2R-MSN activity in females in estrus.^22^ It is possible that drug abuse by males is driven by appetitive drives while in females it is driven by a drive to ameliorate negative affect. It has been suggested that men are more likely to initiate drug use recreationally (i.e. for appetitive reasons) and women are more likely to initiate drug use to cope with negative affect,^37^ which would be behaviorally consistent with the neurophysiological changes observed in animal models. It should be noted that recent work has indicated that in the NAcC the two MSN subtypes work in tandem, with D1R-MSNs responding broadly to stimuli and D2R-MSNs responding to both cues and outcomes, changing with learning, tracking valence-free prediction error, and playing an essential role in associative learning.^38^ Both male and female animals showed locomotor sensitization, confirming that both sexes have behavioral responses to cocaine administration. As no significant cocaine effect was observed in D2R-MSNs or in male D1R-MSNs, and the cocaine effect on D1R-MSNs was estrous-cycle dependent in females, it is possible that the NAcC governs other behavioral responses to cocaine.

The NAc is subdivided into two regions, the NAcSh and NAcC, that have distinct innervation, postsynaptic targets, and physiological roles. It has been suggested that there is a “division of labor” between the NAcSh and NAcC, with the NAcSh thought to regulate reinforcing behaviors such as feeding and drug reward,^39–41^ while the NAcC is thought to regulate learning, motivation, and impulsivity.^12,42–45^ Additionally, the NAcSh and NAcC are thought to play different roles in regulating the response to psychostimulants. The NAcC has been found to play a major role in cocaine-seeking tied to drug-associated conditioned reinforcers, cue-induced reinforcement, and reacquisition after extinction, while the NAcSh regulates the psychostimulant effects of cocaine, such as locomotor sensitization.^12,13^ With these different functional roles in mind, it is unsurprising that regional differences in cocaine-mediated changes in neuroplasticity are observed. The reduction of D1R-MSN excitability following cocaine treatment in the NAcSh^22^ and the absence of cocaine-induced plasticity in the NAcC (Figure 2) suggests that appetitive drive may underlie male drug use more than learned behavior or impulsivity. In contrast, the increased excitability of NAcC (Figure 3) (but not NAcSh)^22^ D1R-MSNs following cocaine exposure in females in diestrus may suggest a role for increased impulsivity in female drug use, especially when estradiol levels are low. The increase in D2R-MSN excitability following cocaine exposure in the NAcSh^22^ and concomitant decrease in D1R-MSN excitability in the NAcC in estrus females (Figure 4) suggests both an increased drive to ameliorate aversive feelings and altered impulsivity may contribute to substance abuse in females. Taken together, there is increasing evidence for an intersection of NAc subregion, sex, and circulating sex hormones in modulating cocaine abuse.

A major unknown, however, is what drives the fundamental sex differences observed in both the NAcC and NAcSh. As fluctuating estrogen levels have been shown to modulate multiple forms of motivated behavior, estrogen signaling seems to be a likely target.^20,46^ As there are two primary estrogen signaling pathways, the nuclear receptor pathway that directly binds to DNA to alter expression and the membrane receptor that activates intracellular signaling cascades via interactions with metabotropic glutamate receptors.^47^ A recent report has indicated that the membrane estrogen receptor modulates alcohol drinking, but not anxiety behaviors, in female mice, highlighting the potential role for specific estrogen signaling pathways in rewarding behavior.^48^ Future studies may look further into this, especially in transgenic models that could assess the role estrogen might play in regional-, sex-, and MSN subtype-specific alterations to neuronal excitability following cocaine exposure.

## Acknowledgements

We would like to thank Dr. Erin Lind and Orion Rainwater of the UMN mouse behavior core for their help, and Dr. Sidney P. Kuo, Chinonso A. Nwakama, and Dr. Andrea R. Collins for proofreading and helpful suggestions.

## Author Contributions

A.D.C., H.M.M., C.H.P., and P.P.J. performed experiments; A.D.C. and H.M.M. analyzed data; A.D.C., H.M.M., and P.G.M., prepared figures; A.D.C., H.M.M., and P.G.M., drafted manuscript; A.D.C., H.M.M., and P.G.M. interpreted results of experiment; A.D.C., H.M.M., C.H.P., P.P.J., and P.G.M. edited and revised the manuscript.

